# A novel tool for optogenetic control of Islets of Langerhans: The Optopus Box

**DOI:** 10.1101/2025.03.18.643880

**Authors:** Samual Acreman, Anton Arturén, Anton Hagsér, Axel Hermanson, Victor Huke, Haiqiang Dou, Johan Tolö, Caroline Miranda

## Abstract

The pancreatic islet hormones insulin, glucagon and somatostatin play a crucial role in maintaining normoglycaemia. Effective communication between beta, alpha, and delta cells is essential for glucose homeostasis and disruption of this intra-islet crosstalk is observed diabetes. Optogenetics is a great way to study pancreatic islets as it eliminates the need for drugs that might have off-target effects, while preserving islet architecture. We generated mouse models of optogenetics to interrogate the physiological and humoral response to electrical and paracrine stimulus in pancreatic islets. To better employ optogenetics on islets, we designed a box to illuminate the islets in batch incubations. Our innovative box is a useful tool for studying islets cell crosstalk in a drug-free, controlled environment and provides a new standard for investigating the dynamic interactions within islets.

## Introduction

The pancreatic islet hormones insulin, glucagon and somatostatin are secreted from β-, α-, and δ-cells respectively^1^ and act to maintain normoglycaemia. Communication between these cells is critical to upholding glucose homeostasis, occurring via paracrine and electrical signals, and disruption of cell-to-cell crosstalk is a key feature of diabetes^2,3^.

Islets cells are electrically excitable. They express ATP-sensitive K^+^ (K_ATP_) channels^4^ that close as glucose concentrations rise, depolarizing the cell and initiating the firing of action potentials. This opens membrane-bound voltage-gated calcium channels (VGCC), enabling the influx of calcium, triggering the assembly of the secretion machinery, releasing hormone. This calcium influx can therefore be used as a proxy signal for secretion at different glucose concentrations in all three cell types^5–7^.

In the study of pancreatic islet physiology, static batch incubation is a standard technique that allows interrogation of hormone secretion from islet cells, within a controlled environment. Drugs that manipulate cell membrane potential via the opening/closing of K_ATP_ channels, or calcium channels blockers^8,9^ are often used in conjunction with this technique to isolate specific intracellular pathways. However, these drugs may impact processes beyond their intended targets, with unwanted physiological consequences^10–12^. Moreover, long term exposure to such drugs may lead to further off-target effects, disrupting metabolic processes and overall cellular health^13^.

During recent decades, optogenetics has been employed broadly across neuroscience^14–16^. More recently, the technology has been applied, with great success, to study islet cells^17,18^. The algal ion channel Channelrhodopsin-2 (ChR2) opens when exposed to blue light^19,20^ at 475nm, and the chloride pump Halorhodopsin (NpHR) which opens when exposed to yellow light at ~590nm^20^can be employed to achieve spatiotemporal isolation of specific cell-types within the islet, while preserving islet architecture, and eliminating the need for drugs ^17,21^. In order to combine batch incubations with optogenetics in a more effective manner, we designed a light box device, named the Optopus Box, tailored for optogenetic stimulation of islets of Langerhans. The Optopus box contains 18 individual tube slots, each equipped with a dedicated blue LED positioned at the base to enable precise illumination and is engineered for complete light isolation. This allows us to combine static batch incubations with optogenetic stimulation, maximising experimental efficacy. Our innovative box, combined with this powerful technology, represents a gold standard for studying spatial and temporal crosstalk between islet cells in a drug-free and controlled environment.

## Materials & Methods

### Selection of Components

Arduino UNO microcontroller was selected for versatility and use friendly properties. For the light source, C503B-BAN-CX0B0461 Cree LED Standard LEDs were selected due to their brightness and specific emission wavelength (470nm). An Adafruit rotary encoder (elektrokit cat#41015725) was used for input control and black 3D printing filament (add:north E-PLA filament) used to print the box.

### Box Design & Manufacturing

The box’s enclosure was designed using Autodesk® Fusion 360 CAD software, incorporating slot dimensions tailored to 3,5mL round bottom polypropylene test tubes. After finalizing the design, the enclosure was printed using an Original Prusa i3 MK3S+ 3D printer.

### Software Development

The control software was developed using Microsoft® Visual Studio Code. The code, written in C, was designed to control the LED brightness, LED frequency and respond to inputs from the rotary encoder. After development, the code was uploaded to the Arduino UNO using Arduino IDE with a USB type a/b.

### Testing

Functional tests were conducted to ensure that the LEDs responded correctly to inputs from the rotary encoder and that the overall system performed as intended. Measurement of illuminance was also conducted to establish the standard deviation of maximal output from each LED using a PM100D-S120C light sensor (Thorlabs).

### Animal Experiments

All animal experiments were performed in accordance with the ethics committee at the Sahlgrenska Academy, at Gothenburg University. Ethical application number 6247/24.

### Optogenetics mouse

We generated mice that harbour the algal protein channelrhodopsin (H134R) under the Lox-Cre system. By crossing Loxed ChR2-YFP ^+/+^ mice (JAX #024109)^22^ with mice harbouring the improved Cre recombinase as described in^23^, under the proglucagon (GCG) promoter, we obtained mice that expressed the ChR2-YFP specifically on α cells.

### Islet Isolation

GCG-ChR2.YFP ^+/+^ and wild type mice were anesthetized and sacrificed by cervical dislocation. Islets were isolated by canulation of the bile duct and injection of 2mL of liberase TL (Liberase Flex, Roche, Basel, Switzerland) diluted at 150ug/ml in Hanks’ Balanced Salt Solution (HANKS’) buffer. Pancreas was digested in a water bath at 37º + shaking (70 rpm) for 8-10 minutes. Digest underwent five washes with HANKS’ buffer supplemented with 5mM glucose + 0.1% bovine serum albumin (BSA;Sigma cat#A7906), before picking of isolated islets from exocrine tissue under a stereo microscope using a pipette.

### Static Hormone Secretion

Islets were preincubated in KRB buffer (0.1% BSA) containing (in mM): 140 NaCl, 4.7 KCl, 2.5 CaCl_2_, 1.1 KH_2_PO_4_, 1.2 MgSO_4_, 25 NaHCO_3_, 10 HEPES (pH was adjusted to 7.4 with NaOH) and supplemented with 6 mM glucose for 1h. They were subsequently transferred to 3.5mL round bottom polypropylene test tubes in groups of 10 islets per tube. Islets were size/mass matched. Each tube was carefully placed in one slot of either the control or the treatment Optopus box, covered with the lid and placed into a shaking incubator, at 100 rpm. The control box was identical to the treatment box, except with the lights turned off. Once both boxes containing the tubes were placed in a cell culture incubator set to 37°C, 5% CO_2_ and the treatment Octopus box light was turned on (delivering continuous illumination). The islets were incubated for 1h, after which, the lights were turned off, the rack was placed on ice, and 150 µl supernatant was collected from each tube and added to collection tubes containing protease inhibitor (Complete, Roche cat# 11697498001) as per the fabricant’s instructions. The samples were stored at −80°C until hormone measurements were performed.

### Hormone Measurement

Glucagon was measured by ELISA (Mercodia, Uppsalla, Sweden).

### Electrophysiology

Membrane potential was recorded with an EPC-10 USB amplifier and PatchMaster software (HEKA Electronic, 658 Germany). Perforated patch technique was employed in the recordings, adding 240 μg/mL 659 amphotericin B to the intracellular solutions to achieve perforation^24^. The extracellular solution contained (in mM): 140 NaCl, 3.6 KCl, 1.3 CaCl_2_, 0.5 MgSO_4_, 10 HEPES, 5 NaHCO_3_, 0.5 NaH_2_PO_4_ (pH adjusted to 7.4 with NaOH), and glucose, as indicated. The pipette solution contained (in mM): 76 K_2_SO_4_, 10 KCl, 10 NaCl, 1 MgCl_2_, 5 HEPES (pH adjusted to 7.35 with KOH). The recordings were performed from islet cells in intact islets held by a glass pipette^25^. Blue light stimulation of channelrhodopsin-2 was applied via LEDs with 470 nm wavelength (Mightex, USA), respectively, from a 60x/1.00 W objective in an upright microscope (Olympus, Japan).

## Results

### The Optopus Box

We developed and manufactured a box for the study of hormone secretion from islets of Langerhans in static batch incubation while under optogenetic stimulation.

The entire box consists of three pieces (Figure 1A-E): a base, a rack, and a lid. The base houses the LEDs and the electronics (Figure 2A-C). The rack has 18 slots for test tubes and a cutout at the bottom of each slot allows 3 mm of the tubes to protrude, for maximum proximity to the stimulating LED on the base of the box. The lid isolates the top of each tube within the confines of a protruding ridge, eliminating light leakage without creating an air-seal. Each piece was 3D printed using black E-PLA filament.

**Figure 1:**
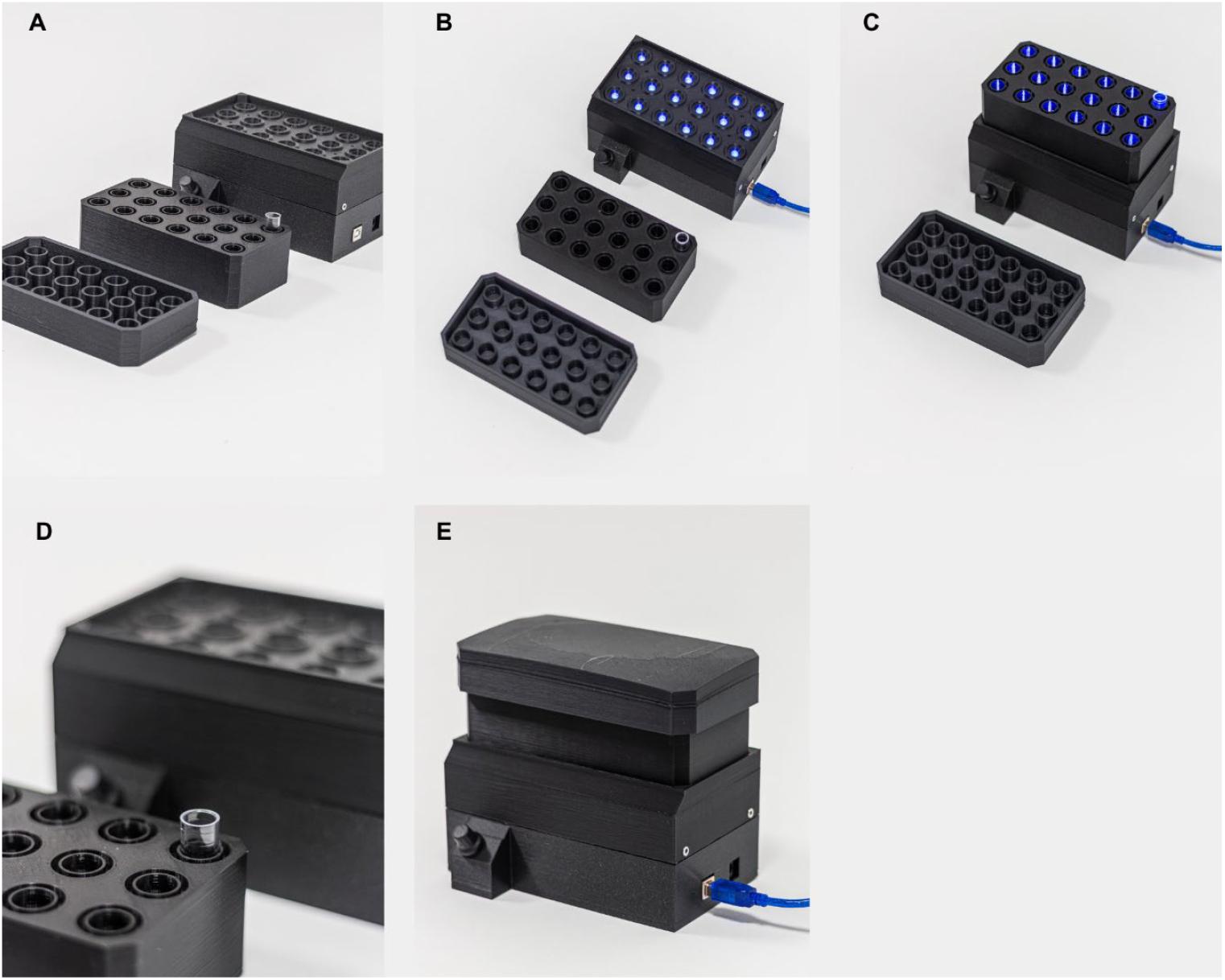
The Optopus Box. 3-D printed in black plastic. It is ultra-low cost (40-50€) and can be adapted to house LEDs of any wavelength for optogenetic studies. The box has a 3-part structure: The base houses 18 LED’s (480nm); the rack for placement of test tubes (tailored for 3,5ml polypropylene tubes); and the lid, which is adapted to individually isolate the tubes (Figure 1 A-E). The dial adjusts the LED parameters can be pressed to swap between continuous illumination and blinking modes.

**Figure 2:**
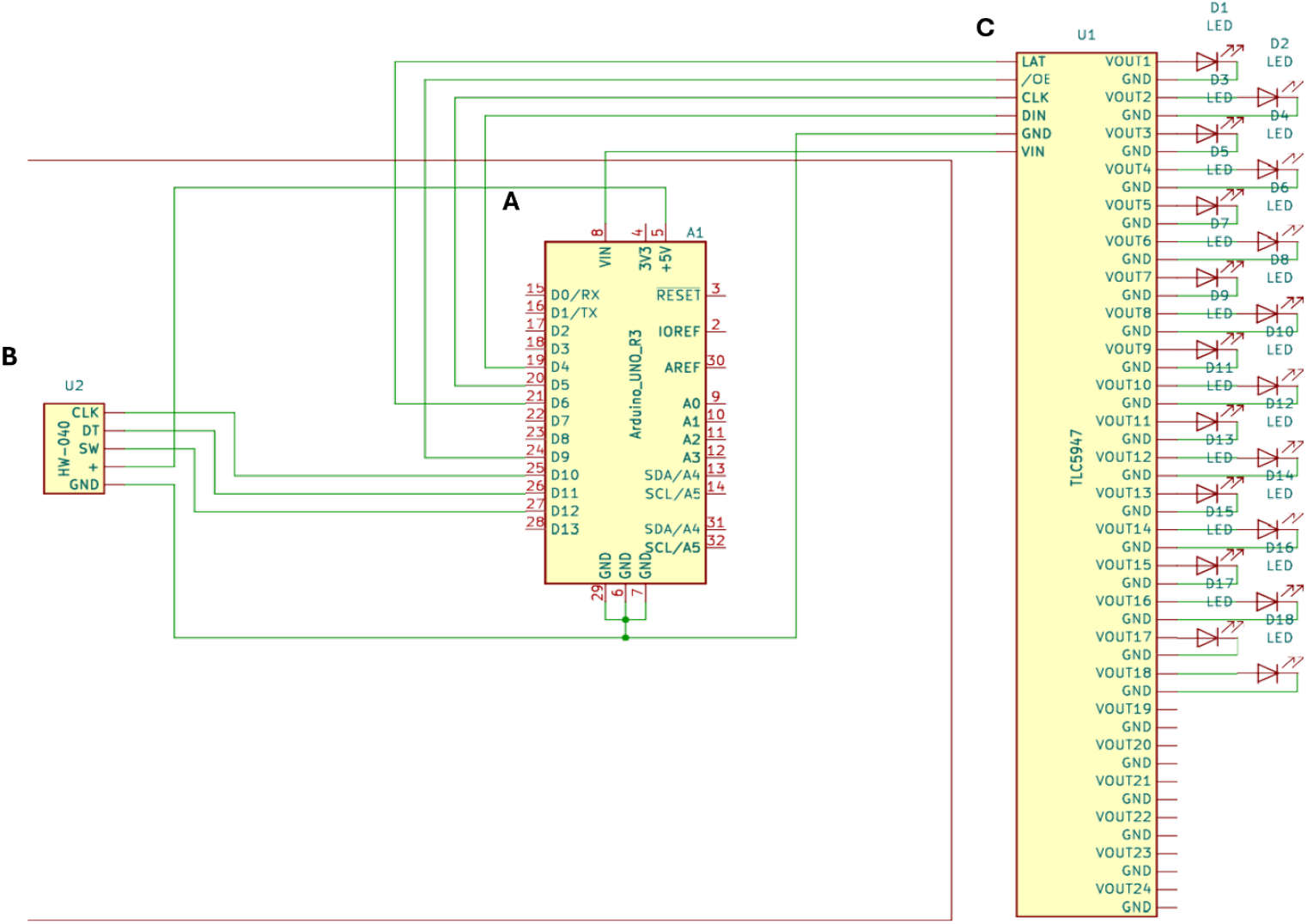
Internal circuit of the Optopus box. **A**: Central square. UNO-R3-Arduino microcontroller houses code that will perform the programmed functions. **B**: Left square. HW-040 - The integrated circuit responsible for generating the CLOCK pulse that determines the operating speed of the ARDUINO. **C**: Square to the right. TLC5947 - the LED Driver that will controls the LEDs (on/off and intensity) based on the commands sent by the Arduino.

### Performance Testing and Validation

The black E-PLA filament was chosen for its colour and density to isolate the tubes from external light, ensuring a reliable test environment. All the software written for this study along with the blueprints for the box are available upon request.

We conducted basic performance tests on the box to ensure its functionality. These included: visual tests of the LEDs response to the rotary encoder: synchronization between the 18 LEDs, increase/decrease of light intensity and blinking interval.

The rotary encoder has an additional button function. Pressing this button switches rotary function from altering light intensity to blinking interval. Light intensity can be increased with clockwise rotation and decreased with counterclockwise. Blink interval can be increased with clockwise rotary rotation whereas counterclockwise decreases the interval between blinks.

Once we asserted that all the above parameters were functional, we measured light intensity/irradiance from all 18 LEDs using a PM100D-S120C light sensor (Thorlabs). All 18 LEDs produced similar irradiance (shown in mW/cm^2^) (Fig 3A). When the rotary was set to 100% output, the average irradiance was 9.67 (±0.14 SEM).

**Figure 3:**
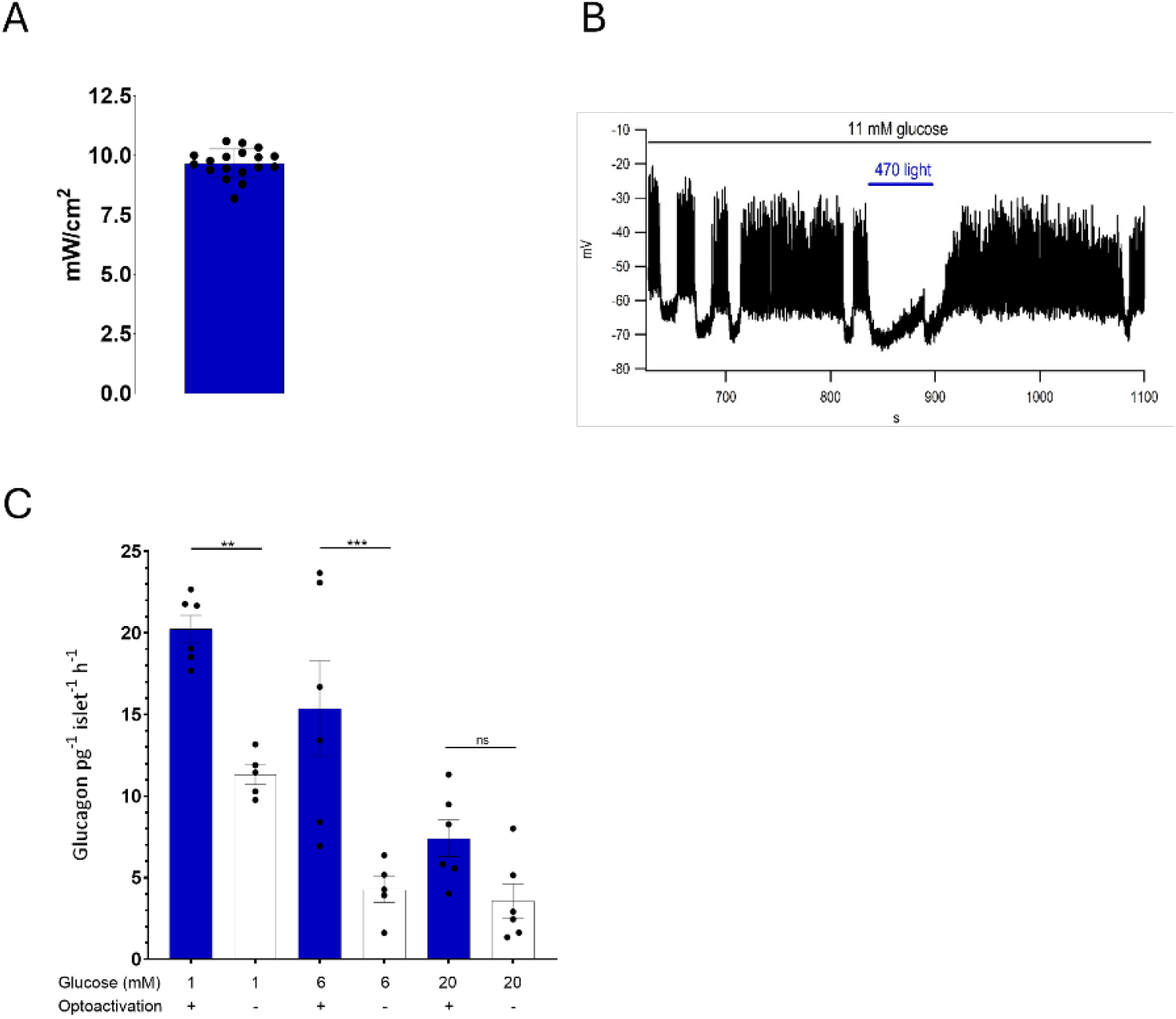
Representative biological experiments conducted for characterization of the optogenetics models, and hormone secretion studies under optogenetic stimulation. **A:** Maximal intensity of the LED’s in mW/cm2. Error bar is ± STD. B: Electrophysiology. Trace from a representative α-cell from within an intact wild-type islet in voltage clamp, where the islet has been exposed to a blue light. The light intensity is set to 100%. Electric activity ceases, due to light toxicity, in the presence of the blue light and resumes when the stimulus is removed. **C:** Representative glucagon secretion from GluiCre islets with/without optostimulation during static incubations. Groups of 10 islets (from 2 mice) were incubated at 1, 6 and 20 mM glucose with, and without incident blue light. Blue bars are lights “on” and white bars are lights “off”. P-values are 0.0041 for 1 mM ON vs 1 mM OFF, 0.0003 for 6 mM ON vs 6 mM OFF, and 0.45 for 20 mM ON vs 20 mM OFF. Error bars show ±S.E.M where statistical significance was calculated using one-way ANOVA (p<0.05 was considered statistically significant).

### Testing for LED cytotoxicity

In the patch clamp setup, the LED delivered a maximum of approximately 3 mW/cm^2^ (Fig 3A). We tested the blue LED on alpha cells from wild-type islets to assess potential adverse effects, a known limitation of optogenetics. At 100% intensity, the blue light caused active cells to cease electrical activity (Fig 3B), which resumed after the stimulus was removed. When tested at 10% intensity, no decrease in electrical activity was observed in wild-type islets. Additionally, in ChR2-Glu islets, a 10% intensity was sufficient to achieve membrane depolarization without harmful effects (unpublished data). Therefore, to avoid any negative impact, we used a light intensity of approximately 10% (~0.3 mW/cm^2^) of the maximum output of the box (equivalent to three steps of the dial).

### Optogenetic control of hormone secretion in mouse islets

In ChR2-Glu islets, glucagon secretion at 1 mM glucose increased from 11.3 pg-^1^ islet-^1^h-^1^ (±0.6 pg) without light stimulation to 20.2 (±0,84pg) in islets that were incubated under incident light (p=0.004). At 6 mM glucose, glucagon secretion increased from 4.27 (±0.8 pg) to 15.4 pg-1 islet-1h-^1^ (±2.9 pg) (p=0.0003). At 20 mM glucose, glucagon increased from 3.6 (±1.0 pg) to 7,4 (±1.1 pg) (p=ns) (Fig 3C).

## Discussion

Normoglycaemia is maintained by the contrasting actions of insulin and glucagon, the two main glucoregulatory hormones, as well as somatostatin, a paracrine regulator of both hormones. Although a consensus on the regulation of β-cell insulin secretion by glucose metabolism has been established, via closure of the K_ATP_ channels and initiation of electrical activity^26–28^, the precise mechanisms through which glucose regulates glucagon secretion remain unclear^7,29,30^. Moreover, the effects exerted by δ-cells on both α- and β-cells, electric or paracrine, have yet to be fully explored^21,31^.

Here we have used optogenetics to stimulate α-cell electrical activity, inducing glucagon secretion in intact islets under both hypoglycaemic and hyperglycaemic conditions. This approach allows investigation of the normal regulation of hormone secretion from the islets of Langerhans, with improved precision and utility with our Optopus box.

### Implications of findings

In this study, we present a key experiment to demonstrate the utility of optogenetics as a means of stimulating pancreatic islets in a clean manner, reducing off-target effects and allowing for precise and rapid (millisecond scale) control over cellular activity, while maintaining the integrity of the islets. While this paper summarizes the crucial aspect of our findings, it is important to note that additional experiments supporting and expanding upon these results are being published in high-impact journals^18^. These comprehensive studies provide a broader context and further demonstrate the utility of the approach documented here.

The box featured in this study was designed and constructed to stimulate islets harbouring the genetically encoded cation channel channelrhodopsin-2^19^, which opens in response to blue light stimulation. Hence, we used LEDs that emit blue light at 470 nm. Furthermore, we have, since then, generated the NpHr mouse model, where islet cells harbour the chloride pump halorhodopsin^20^ (NpHr 3.0), and produced another box, using yellow LEDs. Some of the results obtained from those experiments are published elsewhere^18^.

### Comparison with previous studies

The Optopus box was developed to enhance experimental efficiency and logistical management. Previously, we used an artisanal setup consisting of a plastic box with wires soldered to LEDs, each glued to a tube lid. This required manual attachment to up to 36 tubes to their lids per experiment. This setup was problematic when the racks were placed in the shaking incubator, as the lids would come off the tubes and the soldering would break off partially, causing bad contact, or completely, shutting the LEDs off. The Optopus box significantly improves upon this by streamlining the setup process, eliminating the above problems of shaking-induced damage, thereby reducing manual labour and hence, experiment time.

There are currently, only a handful of studies that use optogenetics to study hormone secretion from pancreatic islets. They have been mainly from our group and using islets from a model of ChR2-RIP, where channelrhodopsin was specifically expressed in mouse β-cells^17,21^. Comparing the results from experiments conducted in the Optopus Box with those from the artisanal setup (unpublished data), the Optopus Box results aligned well with previous experiments, demonstrating its benefits in streamlining the process and reducing setup time. Ultimately, the Optopus Box proves to be the outperformer.

Optogenetics has proven a formidable way to study pancreatic islet physiology. As static incubation is a standard method within the field, optogenetics adds the ability to observe and manipulate cellular processes with high spatiotemporal precision, enabling detailed studies of hormone secretion, cell-cell communication, and intracellular signalling dynamics. In our studies, we used optogenetics to interrogate hormone secretion from islets in static incubation and total pancreas perfusion preparations, as well as electrical activity, and intracellular calcium. All of these, form an optogenetic “toolkit”, that allows for a more in-depth dissection of islet function, advancing our understanding and opening new roads for research and therapeutic interventions.

### Limitations and Technical improvements

Halorhodopsin, a chloride-specific ion pump activated by yellow light, pumps Cl^-^ from the extracellular to the intracellular medium within seconds of activation, independent of the electrochemical gradient^20,32^. However, the biggest caveat with employing this protein to inhibit hormone-secreting cells, is that its “active” state reverts to an inactive state within seconds of initial light activation^32,33^. This inactive state cannot be readily reactivated and requires thermal reversion of the retinal molecule to all-trans-retinoic acid. However, it has been suggested that this can be achieved faster using blue light^32,34,35^. In our studies with the halorhodopsin islets, we have achieved marginal differences in hormone secretion between light-stimulated and unstimulated groups, measured at the end of a 1h incubation (unpublished). With this in mind, we are currently creating a box that possesses both yellow and blue LEDs at the base, for double illumination.

The new box has a more advanced software to control the processor. It can be accessed and edited from a computer and allows us to program diverse timing intervals between “lights on” and “lights off” across both wavelengths.

Despite the success of the Optopus box in decreasing the preparation time for experiments, there remains room for improvement. The primary improvement would be to the power supply for the box. The shaker-incubator where we perform our experiments is made of Plexi glass and has a small hatch under the main door. This hatch can be removed to reveal an opening through which we pass the cable. The incubator has temperature control, so keeping the temperature during the 1h is feasible. However, making the box battery driven, would improve its portability and eliminate the need for external power sources. Other improvements we are developing are the use of PET plastics to print the box, a more user-friendly (easier to adjust blinking interval and light intensity) interface. In implementing these improvements, we hope to further simplify the setup process, and to increase the applications of the box (other tissues, other fields).

## Acknowledgements

CM had a fellowship from SSMF (Sweden) at the time of the study.

Edmar Miranda and Rickard Larsen, for technical consulting. NTI Johanneberg for covering the costs associated with production of the boxes.

## Author Contributions

SA conceptualized and designed the initial prototype. AA, AH and AH made the blueprints, 3-D printed the parts, wrote the code and assembled the box. VH supervised the manufacturing of the box as main engineer. CM, JT, and HD conceptualized the final version of the box, designed and performed the biological experiments, analysed the data, discussed and designed the manuscript. CM wrote the paper. All authors discussed the manuscript and approved its final version.

